# Low *P*_crit_ but no hypoxia tolerance? Hypoxia compensation in the Arctic keystone species *Boreogadus saida*

**DOI:** 10.1101/2023.05.04.539365

**Authors:** Sarah Kempf, Carolin Julie Neven, Felix Christopher Mark

**Affiliations:** Integrative Ecophysiology, Alfred Wegener Institute Helmholtz Centre for Polar and Marine Research, Am Handelshafen 12, 27570 Bremerhaven, Germany; University of Bremen, Bibliothekstraße 1, 28359 Bremen, Germany; Channel and North Sea Fisheries Research Unit, Ifremer, Boulogne-sur-Mer, France

**Keywords:** hypoxia, ocean warming, metabolic scope, *P*crit, swimming performance, Polar cod

## Abstract

Global warming has already caused a loss of almost 50% Arctic sea-ice coverage since the 1980s. Sea-ice loss strengthens summer stratification of the ocean’s water column and, consequently, hypoxic zones in the deep-water layers may form. The present study investigated the response of an Arctic keystone species, the Polar cod, *Boreogadus saida*, to hypoxia and warming. We measured the respiratory capacity (standard, routine and maximum metabolic rates, SMR, RMR, MMR, aerobic scope, critical oxygen saturation (*P*_crit_)) and swimming performance of Polar cod under progressive hypoxia at 2.4 °C and after warm acclimation to close to the species’ thermal limit (10.0 °C) via flow-through and swim tunnel respirometry. We observed clear and stable patterns that were similar in both thermal regimes: Polar cod displayed oxygen-regulating behaviour under progressive hypoxia, with SMR never below aerobic baseline metabolism and a very stable AS. Our study revealed that Polar cod can handle exceptionally low oxygen saturations down to a *P*^crit^ of 5.9 % air saturation at typical habitat temperatures. Closer to critical temperatures (10.0 °C), *P*_crit_ rose to 21.6 % air saturation. However, the pertinent question remains whether the observed behaviour can be summarized under classic hypoxia tolerance, as we a) did not observe any metabolic downregulation and b) no anaerobic component of the hypoxia response in Polar cod, which are usually put forward in the definition of hypoxia tolerance. Therefore, we describe the observed metabolic response to hypoxia rather as metabolic hypoxia compensation than hypoxia tolerance as the mechanisms involved here actively seek to improve oxygen supply instead of (anaerobically) tolerating hypoxia through metabolic depression.

## 1 Introduction

Arctic ecosystems are characterized by strong seasonal fluctuations in sea ice coverage and formation. Currently on-going climate change is leading to rapid warming of the Arctic, and due to Arctic amplification, the Arctic oceans are expected to warm twice as fast as the global average [1, 2]. Due to increased sea surface temperatures, the Arctic has already lost 49 % of its sea ice, compared to the 1979-2000 baseline of 7.0 x 10^6^ km^2^ [3, 4] and according to a business-as-usual greenhouse gas emission scenario (RCP8.5) [5-7], it is projected that the Arctic will be nearly ice-free in September before the year 2050 [8].

This dramatic sea ice loss has also been observed in the fjord systems of the Svalbard archipelago [9, 10]. While some of the Svalbard fjords are widely open to the neighbouring sea, others are not and can be regarded as closed, or semi-closed, due to their relatively shallow sills (e.g.: van Mijenfjorden, Billefjorden, Brepollen/Hornsund) [10-12]. In these systems, loss of sea ice and declining sea surface salinities due to freshwater inflow can lead to increased upper ocean stratification. In stagnant deep-water bodies, this will result in O_2_ depletion by biological processes, which can pose a challenge for marine organisms.

In semi-closed fjord systems, the thermohaline circulation caused by sea ice formation is the main driver of the abiotic conditions in the deep-water layers [13]. The seasonal downward transport of cold, dense and highly saline water from the surface to the bottom breaks up summer stratification and thereby replenishes oxygen and nutrient levels in the bottom waters with locally formed winter cooled water (WCW) [12, 14]. The reduced ice formation or even a complete lack of ice cover is having serious ecological consequences as the deep cold-water layers are not receiving sufficient amounts of oxygen-rich water and oxygen depletion may extend over more than one season [15, 16] . The replacement of WCW by local- and intermediate water in warmer years, as demonstrated for Hornsund in Svalbard [12], can potentially lead to local bottom hypoxia.

In Billefjorden oxygen levels regularly decrease from 100% air saturation (366.67 µmol O_2_/l) in top water layers down to 75% air saturation (275.01 µmol O_2_/l) at the bottom at the end of summer (Figure 1A).

**Figure 1.**
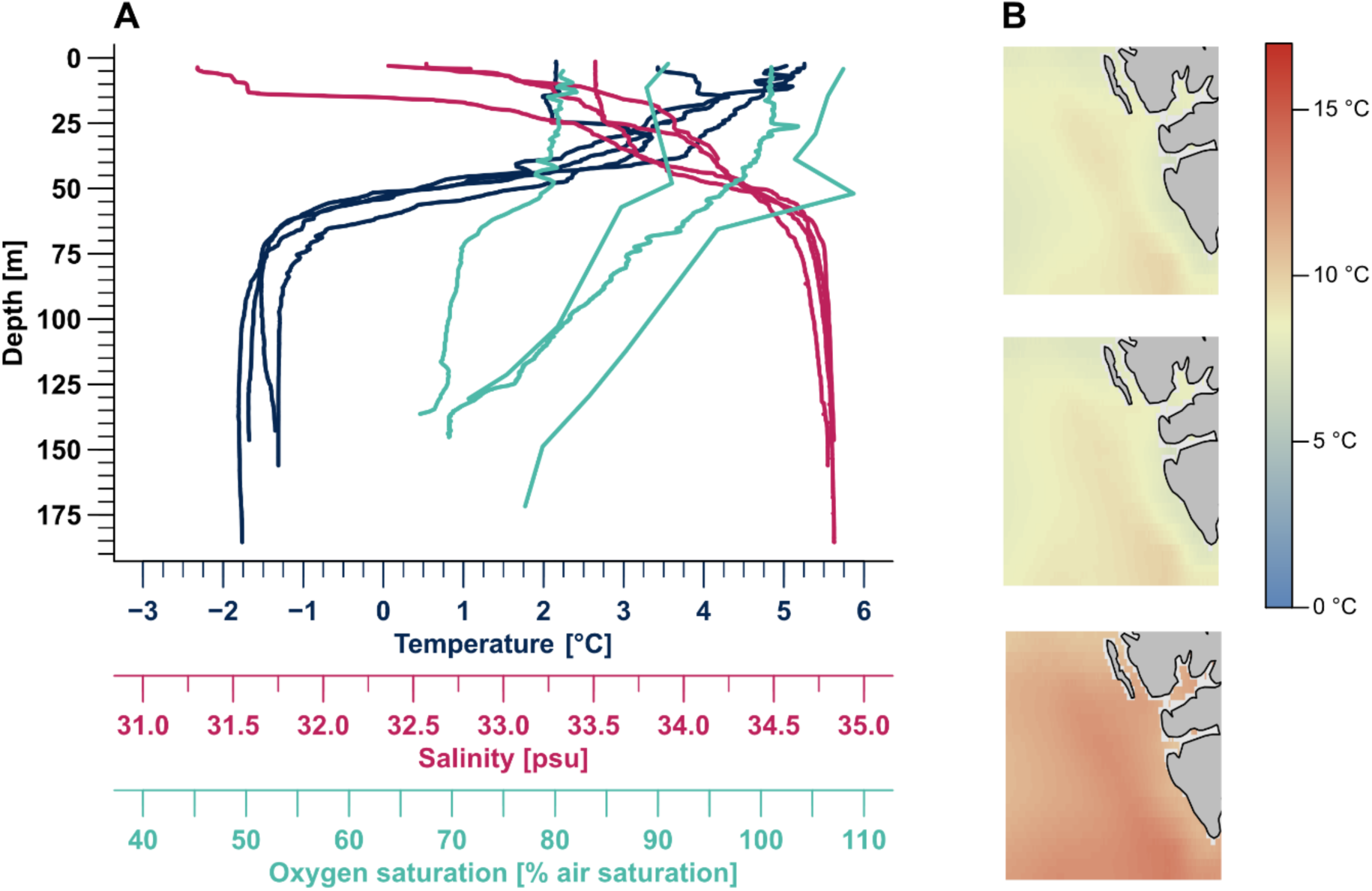
CTD profiles (2013 – 2020) and sea surface temperature models for 2100 of the study area. **A:** Vertical temperature – (dark blue, in °C), salinity – (pink, in psu), and oxygen profile (turquoise, in % air saturation) of Billefjorden. CTD profiles casts were performed in the years 2013, 2015, 2018, 2020 with the RV Heincke (data from: https://doi.org/10.1594/PANGAEA.824703; https://doi.org/10.1594/PANGAEA.855528; https://doi.org/10.1594/PANGAEA.896218; https://doi.org/10.1594/PANGAEA.928667). Due to technical problems with the oxygen sensor, we were only able to record a small number of oxygen saturations during the CTD cast in 2015. In 2020, the oxygen sensor could not be delivered in time due to the corona pandemic; the oxygen saturation was measured manually after the CTD cast using an optode. **B:** Calculated sea surface temperature for the year 2100 along the west coast of Svalbard after RCP 4.5 (first), RCP 6.0 (second) and RCP 8.5 (third) [5] (modelled in R, BioOracle).

One of the most abundant fish species inhabiting these deep fjord basins is Polar cod, *Boreogadus saida.* Feeding on small invertebrates such as copepods, amphipods, gastropods and krill, and being themselves a main food source for large fish, marine birds, and mammals, Polar cod build the link between the lower trophic levels and top predators, and are therefore considered a key species in the Arctic ecosystem [17]. Dependent on life stage, *B. saida* is found in both pelagic and benthic habitats. While the young of the year occur mostly in relatively warm surface and midwater layers (up to 6°C, pers. obs. FC Mark; Figure 1A) during their first summer, juveniles are often ice associated. Adult individuals spend the winter seasons in both pelagic and bottom waters and retreat to cold WCW layers at the fjord bottom in summer (cf. Geoffroy et al., in final review for Elementa, MS No. ELEMENTA-D-22-00097, awaiting acceptance soon). These isolated fjord basins are protected from warming and, due to their very low temperatures (down to -1.9 °C, Figure 1A) also from predation pressure by invading species from Atlantic ecosystems such as Atlantic cod, *Gadus morhua* [18-21].

Due to their key role in the Arctic food chain, the ecology, physiology, and resilience of *B. saida* to climate change has been the focus of recent studies [e.g. 17, 22, 23-28], and it has since become a model species. In the context of the ongoing transformation (and Atlantification [29]) of the Arctic ecosystem around Svalbard and possible future deoxygenation of its deep fjords, we here assessed the hypoxia and thermal tolerance of metabolic performance of *B. saida*. Specifically, hypoxia tolerance has never been investigated in Polar cod – and generally very scarcely in polar fish – partly owing to the paradigm that polar ectotherms have never evolved hypoxia tolerance as they are normally not exposed to oxygen-poor conditions in the cold polar waters [30-33]. To close this gap in knowledge, and predict future climate change scenarios in these already strongly affected ecosystems, we need to understand the physiological response of this Arctic key species to local hypoxia and rising temperatures [34-36].

In the present study, we assessed hypoxia tolerance and metabolic performance under progressive hypoxia at optimum temperature (2.4 ± 0.6 °C) and after long-term (12 months) acclimation to 10°C, which is close to Polar cod’s critical temperature T_crit_ of 12°C. We focused on the standard (SMR), active (AMR), maximum metabolic rate (MMR) and aerobic scope (AS) to evaluate how progressive hypoxia influences *B. saida*’s metabolic rate, and to identify its critical oxygen concentration (*P*_crit_). We further analysed their swimming performance to determine *P*O_2_ dependent U_gait_, the speed at which the fish transitioned from steady aerobic to an anaerobically supplemented “kick-and-glide” swimming gait, and the maximum swimming velocity U_crit_ achieved in the swim trials [cf. 23].

## Materials and methods

Polar cod for the experiments in this study were caught in October 2018 during RV Heincke expedition HE519 in Billefjorden (78°34’59.99" N 16°27’59.99" E), the innermost part of the Isfjorden, which is located at the west coast of the Svalbard archipelago (Figure 1). The fish were caught with a fish-lift connected to a pelagic trawl [37] at 150 m depth, at a temperature of -1.5 °C, and an oxygen level of 75% air saturation (275.01 µmol O_2_ / L). Once transported to the Alfred Wegener Institute (AWI) in Bremerhaven, Germany, fish were maintained in two flow-through tanks at normoxia and at an ambient temperature of 0.0 ±0.5 °C.

For the experiments, 46 fish were tagged with passive glass transponders (PIT tag, FDX-B, 7 x 1.35mm, Loligo Systems Denmark) close to the dorsal fin, after anesthetization with tricaine methanesulfonate (MS-222; 0.125 g L^-1^). During this procedure, fish were also measured (total and standard length, width and depth) and weighed. No later mortality due to tagging and handling stress occurred.

These 46 fish were divided into two groups. The first group (n = 30; labelled C, cold) measured 19.7 ± 1.3 cm and 39.6 ± 9.5 g and was kept in the above-mentioned aquarium tanks. The second group (n=16; labelled WA, warm acclimated) with an average size of 19.8 ±1.9 cm and weight of 41.3 ± 10.1 g, was progressively acclimated to 10.0 °C (warming rate: 1.5 °C month^-1^) and then kept at this temperature for several months. The temperature of 10°C is close to the critical temperature of Polar cod (long-term mortality increases sharply beyond 12 °C, pers. obs. FC Mark). Furthermore, 10.0 °C is within the thermal range of the projected future surface water temperatures for the western fjord systems of Svalbard according to RCP 8.5 [5]. Projections were modelled in R language (using the packages “sdmpredictors”, “tidyverse”, “raster”, “sp” “dismo”, and “leaflet”). Models were generated using bathymetry and environmental data derived from Bio-ORACLE at a resolution of 5 arcmin or 9.2 km [https://bio-oracle.org/, 38, 39], for western fjord systems of Svalbard, using the mean sea bottom temperature of 2.4 ± 0.1 °C under RCP 8.5 [5] for the year 2100.

### 2.1 Metabolic rate measurements

For the first experimental group (C), an experimental temperature of 2.4 ± 0.6°C was chosen for respiration chambers and swim tunnel, corresponding to the optimal temperatures for Polar cod as previously identified for metabolic rate, mitochondrial performance and growth [22, 25]. The second group (WA) was measured at their acclimation temperature of 10.8 ± 1.0 °C to gauge the remaining aerobic scope close to T_crit_ (long-term mortality increases sharply beyond 12 °C, pers. obs. FC Mark).

#### 2.1.1 SMR and RMR measurements

Standard (SMR) and routine (RMR) metabolic rates were measured using seven fully automated respiration chambers (Loligo Systems ApS, Denmark), submerged in two connected thermoregulated tanks (170 L). The water from the outer reservoirs was re-circulated to each respirometer using computer-controlled flush pumps (Compact 600, EHEIM, Germany), relays and a software (AutoResp 2.3.0, Loligo Systems ApS, Denmark). Each respirometry chamber also had its own circulation loop including an oxygen sensor (Witrox 4 oxygen meter, Loligo systems ApS, Denmark) that continuously monitored water oxygen level inside the chamber. Oxygen probes were calibrated to 0% air sat (nitrogen saturated water) and 100% air sat (fully aerated water) at the beginning of each respirometry trial.

As soon as fish were introduced in the respirometry chambers, the oxygen consumption (*Ṁ*O_2_) measurement sequence was started. Measurement cycles consisted of an open period, during which respirometers were replenished with water from the surrounding reservoir, and a sealed period, when the flushing pumps were turned off and the decline in water oxygen over time used to calculate the corresponding fish *Ṁ*O_2_. Note that the first 200 s of the measurement period was not used for the calculation of *Ṁ*O_2_ (stabilization period). The respirometers’ measurement cycles were 5 min flush, and 30.5 min measurement in the cold group (C) and 2.5 min flush, and 17.5 min measurement in the WA group. The respirometry set-up was located in a space with restricted access and visual contact between the animals was prohibited by opaque plastic walls between the chambers. Note that fish were fasted 7 days (group C) or 3 days (group WA) prior to being transferred to the respirometers.

Each oxygen saturation was maintained for two days and two nights, containing approximately 80-100 measurement phases during which fish were left undisturbed to ensure proper determination of the standard metabolic rate (SMR) as they habituated to the experimental conditions [40]. Only the metabolic rate data from the second night were used for SMR calculation. Oxygen consumption was measured in 12 different *P*O_2_ conditions, 100, 75, 65, 55, 45, 35, 30, 25, 20, 15, 10 and 5% air saturation (n = 7 per conditions).

The rates of oxygen consumption were calculated after Boutilier et al. [41] and normalized according to Steffensen et al. [42] (see below: 2.3 Data handling) using the statistical software “R” with the packages “FishResp” [43] and “Mclust” [44].

Blank respiration was carried out after each experimental run and subtracted from individual fish respiration. All chambers were cleaned and disinfected weekly.

### 2.2 Swimming performance measurements

The metabolic rate and swimming performance of *B. saida* under hypoxia were recorded following a critical swimming speed (U_crit_) protocol [Brett (1964), modified after 23], applying the same *P*O_2_ steps as in the respiration chambers (100, 75, 65, 55, 45, 35, 30, 25, 20, 15 and 10 % air saturation).

A Brett-type swim tunnel respirometer of 5 L (test section 28 x 7.5 x 7.5 cm, Loligo Systems ApS, Denmark) was used to measure the swimming performance of *B. saida* (n=6-7 per *P*O_2_ treatment). Oxygen content and flush phases were controlled by a DAQ-M instrument (Loligo Systems ApS, Denmark). The swim tunnel was submerged in a reservoir tank to maintain stable abiotic conditions within the chamber. The water velocity was regulated by a control unit regulating the rpm of an engine and connected propeller (Loligo Systems ApS, Denmark). To calibrate the water velocity to voltage output from the control system, a vane wheel flow probe (Loligo Systems ApS, Denmark) was used.

The *P*O_2_ was determined using fiber optic mini sensors (optodes) connected to a four-channel oxygen meter (Witrox 4 oxygen meter for mini sensors, Loligo Systems ApS, Denmark).

The fish were transferred to the swim tunnel, seven days (C) or three (WA) days after the last feeding. After an acclimatisation period of 1.5 h in stagnant water, water velocity was increased to 1.2 BL (body length)/sec for 25 min. Afterwards, the velocity was increased to the first measurement velocity of 1.4 BL/sec before starting the U_crit_ protocol. Each velocity step contained an *Ṁ*O_2_ measurement cycle, comprising a 60 s flush phase followed by 120 s of stabilisation and an 8 min measuring period, after which water velocity was increased by 0.15 BL/sec. Each measurement period thus returned 480 single oxygen recordings from which oxygen consumption (*Ṁ*O_2_) was calculated. The swim chamber was covered to minimise disturbance.

This stepwise increase was performed until exhaustion, when the fish completely refused to swim and remained inactive for more than three minutes. The maximum metabolic rate was determined for each *P*O_2_ level inside the swim tunnel.

To determine the gait-transition speed U_gait_ (the switch from strictly aerobic to anaerobically supplemented swimming) [45], kick-and-glide swimming (so called “bursts”) [46] was documented. In kick-and-glide swimming, thrust generation is supplemented by anaerobic muscle contractions and mainly the white muscles are used. All bursts were counted and the corresponding time during the swim trial was documented. The critical swimming velocities (U_crit_) of the fish were calculated according to Brett [47] (see: 2.3.6). After U_crit_ was reached, the velocity was immediately decreased to the basic weaning velocity of 1.4 BL/sec and the fish stayed in the swim tunnel for another 10 min before being transferred back into their tanks.

Blank respiration in the swim tunnel accounted for less than 2% *Ṁ*O_2_. The swim tunnel was cleaned daily.

### 2.3 Data handling

Oxygen consumption (*Ṁ*O_2_) and the corresponding metabolic rates for both experiments were calculated from optode oxygen recordings measured in the respiration chambers and swim tunnel, and normalized in R language [48] using the package “FishResp” [43]. Afterwards *Ṁ*O_2_ was manually normalized after Steffensen et al. [42] to a body weight (BW) of 100 g using the following equation

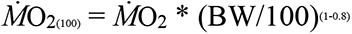

#### 2.3.1 Baseline SMR calculation

The baseline SMR was calculated after Chabot et al. [40], using the function “Mclust” of the R package “mclust” [44]. Briefly, stable *Ṁ*O_2_ between 100 % and 60 % air saturation of all fish were used as a baseline for this calculation. The lowest 5 % of these data were removed as outliers, afterwards the mean value of the remaining lowest 15 % (quantiles) was calculated and used as baseline SMR for subsequent metabolic rate analyses [40].

#### 2.3.2 SMR calculation

Different to baseline SMR, SMR was calculated for each individual from the respective *Ṁ*O_2_ data recorded in the respiration chambers. For each fish within a *P*O_2_ group (5-100 % air saturation), approximately 45 individual *Ṁ*O_2_ data points were calculated, the lowest 5 % of these data were discarded as outliers and the mean of the remaining lowest 15 % was calculated as individual SMR. The mean value of all calculated individual SMRs in a treatment was calculated, and determined as the specific SMR for each treatment. The standard deviation was calculated for each *P*O_2_ group. All calculations were performed in R language using the above-mentioned packages.

#### 2.3.3 MMR calculation

For MMR calculations only, the data obtained in the swim tunnel experiment were used. Due to the short measuring time of maximum two hours, the number of data points per fish (between 3 and 10 individual *Ṁ*O_2_ data points) was too small to calculate the mean value of the highest 15% for each fish after the highest 5% were removed as outliers. Therefore, the highest 5 % of all recorded *Ṁ*O_2_ measurements in each treatment were removed as outliers. The remaining highest metabolic rate per fish was taken as MMR. The mean value of all individual MMRs and the corresponding standard deviation was calculated for each *P*O_2_ group.

#### 2.3.4 Aerobic scope calculation

Based on the individual SMR and MMR data, the AS per *P*O_2_ group was calculated in R using the package “tidyr” [49] after following equation:

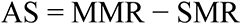

#### 2.3.5 *P*_crit_ calculation

The “level of no excess activity” [50], “*P*_crit_ (P_c_)” [51], or “O_2crit_” [52], the minimum oxygen level required to maintain standard metabolism has various names. All of them describe the oxygen threshold below which fishes switch to anaerobic metabolism, survival depends on how far below SMR the metabolism stabilizes and for how long the situation persists [52, 53]. Below this threshold, oxygen shifts from a limiting factor to a lethal factor. It is calculated from the point of intersection between MMR and baseline SMR, based on their regression analysis with the function “curve_intersect” of the R package “reconPlots” [54].

If oxygen conditions below *P*_crit_ persist for some time, or if the oxygen content continues to decrease, the “incipient lethal oxygen level” (ILO), defined as the loss of equilibrium, is reached [52, 55, 56]. The small range between *P*_crit_ and ILO can be classified as the “scope for survival”, the difference between the SMR and the minimum achievable metabolic rate [57].

#### 2.3.1 *P*_crit-max_ calculation

*P*_crit-max_, the *P*O_2_ at which oxygen supply fails to cover the maximum demand for oxygen resulting in a decrease of MMR [58-60], can be calculated as the intersection of a linear function [α-line,58] with a slope α = MR*_P_*_crit_/*P*_crit_ with MMR.

#### 2.3.2 Critical swimming speed (**U_crit_**) after Brett (1964)

The critical swimming speed was calculated as:

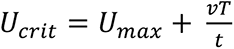

With U_max_ as the highest velocity (v) perpetuated for a complete time interval (t) and T as the time spent at the given velocity leading to exhaustion of the fish.

#### 2.3.3 Gait transition speed (**U_gait_**)

The critical swimming speed was calculated as:

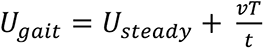

With U_steady_ as the highest velocity (v) perpetuated for a complete time interval (t) without burst swimming and T as the time spent at the given velocity leading to burst swimming.

#### 2.3.4 Statistical analysis

For each experiment, data were tested for normal distribution using either a Shapiro-Wilk test or Kruskal-Wallis test. Homoscedasticity was tested for using Levene’s test. The significance of the differences between *P*O_2_ treatments were evaluated by one-way (respiration chambers, burst counts and active swimming duration) or two-way (swim tunnel) ANOVA followed by Tukey’s test for mean value comparison. Furthermore, the influence of body weight, body length, temperature and water velocity were examined. The level of statistical significance was set to p < 0.05 for all statistical tests. All statistical tests were performed in R language (version: 3.6.1).

## 3 Results

### 3.1 Mortality

During the experiments at 2.4 °C, mortality (n=2) occurred only during the acclimation phase at the lowest *P*O_2_ (5 % air saturation). Another four individuals died in the rearing tank over the course of the experiment, independent of handling stress or *P*O_2_.

At 10.0 °C, two individuals died at 25 % air sat. in the respiration chambers. Therefore, the RMR measurements were prematurely terminated and SMR had to be calculated from the *Ṁ*O_2_ data of the five remaining individuals. During the swim tunnel experiment, one individual died at the end of the acclimation period to 20 % air sat.. Another seven individuals died in the rearing tank over the course of the experiment, independent of handling stress or *P*O_2_.

### 3.2 Respiration measurements

Metabolic rate was significantly influenced by body weight (p_RMR_ < 0.0001, p_MMR_ = 0.0039). Additionally, AMR (active *Ṁ*O_2_) was also found to be significantly influenced (p = 0.0082) by body length. At both temperatures, oxygen consumption was elevated upon the introduction of fish in the respirometry chambers, but progressively declined to a steady level after about 36h. Therefore, we only used *Ṁ*O_2_ data of the second night to calculate routine and standard metabolic rates.

#### 3.2.1 Baseline standard metabolic rate

The baseline SMR of *B. saida* was calculated after Chabot et al. [40] based on the *Ṁ*O_2_ data recorded between 60 and 100 %. At 2.4 °C, it amounted to 0.37 µmol O_2_/g·h. After acclimatisation to 10.0 °C, SMR was significantly increased almost 6-fold, reaching 2.12 µmol O_2_/g·h (p < 0.001).

#### 3.2.2 Routine and standard metabolic rate

At both temperatures, SMR did not decrease below baseline metabolism (baseline SMR) throughout the oxygen range.

At 2.4 °C, *Ṁ*O_2_ was significantly dependent on *P*O_2_ (p < 0.001). SMR (Figure 1 A, Table 1) remained on a steady level between 100 and 45 % air sat. (332 – 155 µmol O_2_/L; max: 0.82 ± 0.69, min: 0.45 ± 0.11 µmol O_2_/g·h), below which oxygen consumption increased between 45 and 35% air sat. (155 – 115 µmol O_2_/L) up to 1.31 ± 0.17 µmol O_2_/g·h (p < 0.001). From 35 to 20% air sat. (115 – 73 µmol O_2_/L) the *Ṁ*O_2_ decreased down to 0.90 ± 0.17 µmol O_2_/g·h (p < 0.0001) and remained stable between 20 and 5% air saturation (73 – 24 µmol O_2_/L).

**Table 1.**
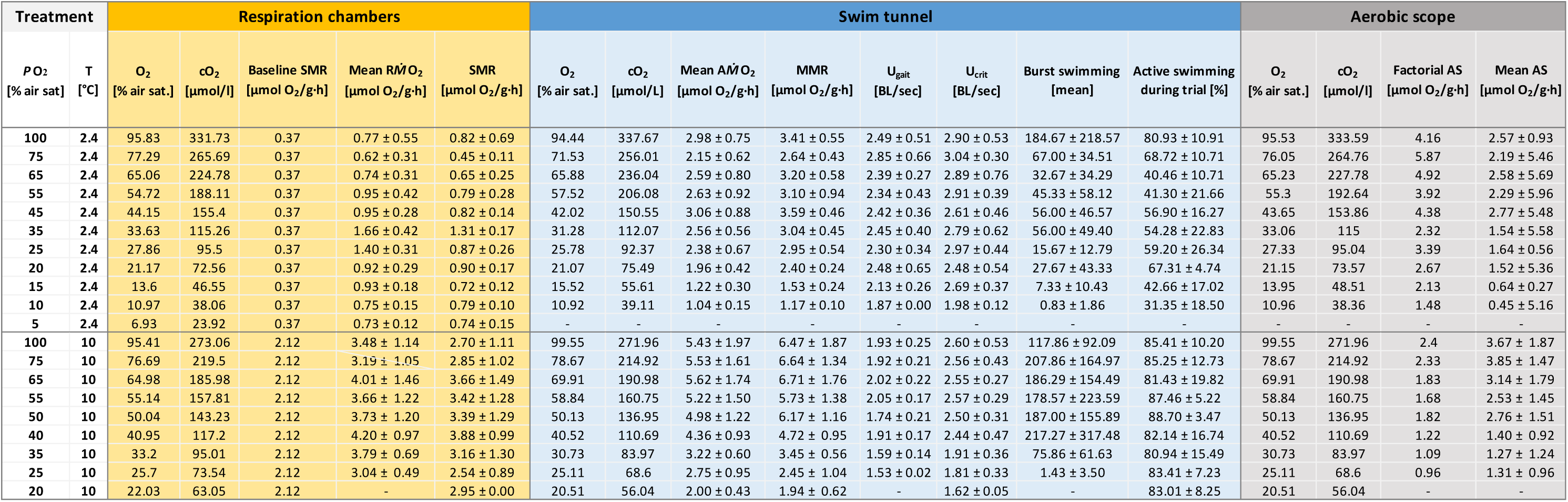
Summary of the main results from both temperature treatments. The first column show the results of the respiration chambers and include the ambient oxygen saturation in % air saturation and µmol O_2_ /L of each oxygen treatment, the baseline SMR, mean R*Ṁ*O_2_ and SMR (in µmol O_2_/g*h). To summarize the swim tunnel experiments, the ambient oxygen saturation in % air saturation and µmol O_2_ /L of each oxygen treatment, the mean A*Ṁ*O_2_ and MMR (in µmol O_2_/g*h), U_gait_ and U_crit_ (in BL/sec) and percentage of active swimming during the trial are listed in the second column. As link between routine and active metabolism, the AS (as mean of each O_2_ treatment, in µmol O_2_/g*h) is summarized in the last column.

SMR at 10.0 °C was not significantly affected by progressive hypoxia, but significantly elevated by temperature (p < 0.001) (Figure 1 B, Table 1). It was maintained between 100 and 55 % air sat. (286 – 186 µmol O_2_/L; 3.0 – 3.5 µmol O_2_/g·h) followed by a slight increase at 40-50 % air sat. (143 – 114 µmol O_2_/L) up to 3.88 ± 0.99 µmol O_2_/g·h which levelled off towards 25% air sat. (71 µmol O_2_/L; 2.54 ± 0.89 µmol O_2_/g·h).

#### 3.2.3 Active and maximum metabolic rate

The *Ṁ*O_2_ during the swim tunnel experiment at 2.4 °C was significantly influenced by *P*O_2_ (p < 0.0001) and water velocity (p < 0.001). MMR (Figure 2 A) was maintained between 100 and 45 % air sat. (3.2 µmol O_2_/g·h), with a small, insignificant decline at 71.53 ± 1.84 % air saturation (256.02 ± 4.71 µmol O_2_/L) down to 2.64 ± 0.64 µmol O_2_/g·h. Beginning with 31.28 ± 1.52 % air saturation (112.07 ± 1.70 µmol O_2_/L) the *Ṁ*O_2_ (3.04 ± 0.45 µmol O_2_/g·h) decreased in parallel to decreasing oxygen content until a metabolic rate of 1.17 ± 0.10 µmol O_2_/g·h was reached at 10.92 ± 1.50 % air saturation (39.12 ± 0.59 µmol O_2_/L).

**Figure 2.**
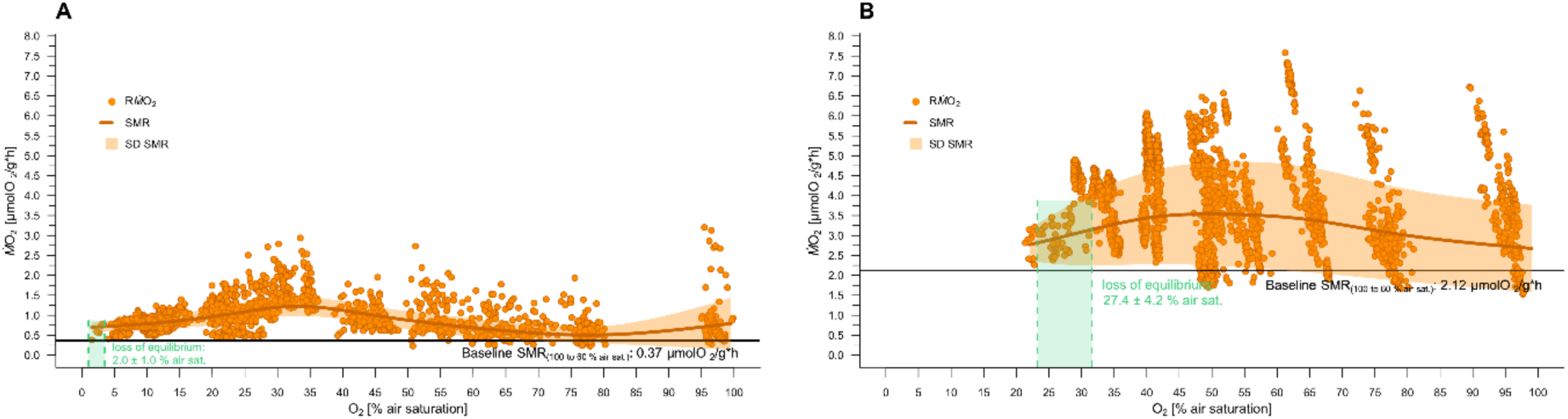
Routine metabolism and standard metabolic rate with progressing hypoxia at 2.4 and 10.0 °C. The oxygen consumption ((*Ṁ*O_2_) in µmol O_2_/g·h) over oxygen saturation (in % air saturation) is displayed as routine oxygen consumption (orange circles, R*Ṁ*O_2_) and standard metabolic rate per treatment group (dark orange line, SMR as polynominal regression) with standard deviation (orange shaded area, SD SMR). At the lower end of the oxygen range, fish lost equilibrium (green shaded area), which terminated the experiment. For both temperature treatments, the baseline SMR has been calculated from RMRs measured between 60 and 100 % air sat. (Baseline SMR). **A:** *Ṁ*O_2_ at 2.4 °C. 1887 single *Ṁ*O_2_ data points representing 86 experimental runs of 30 individuals (7 individuals per treatment). The fish lost equilibrium at 2.0 ± 1.0 % air sat. (7.0 ± 3.5 µmol O_2_ /l). The baseline SMR accounted for 0.37 µmol O_2_/g·h. **B:** *Ṁ*O_2_ at 10.0 °C. 2473 single *Ṁ*O_2_ data points representing 53 experimental runs of 23 individuals (7 individuals per treatment). The fish lost equilibrium at 27.4 ± 4.2 % air sat. (78.5 ± 3.3 µmol O_2_ /l): n=5 for 25% air sat.. The baseline SMR accounted for 2.12 µmol O_2_/g·h.

At 10.0 °C and full oxygen saturation, MMR was significantly higher compared to 2.4 °C (6.71 vs. 3.2 µmol O_2_/g·h, respectively. p < 0.001). Progressive hypoxia had a significant negative effect on MMR (p < 0.001). Between 100 and 70 % air sat. (274 – 205 µmol O_2_/L), *Ṁ*O_2_ was maintained (6.71 ± 1.76 µmol O_2_/g·h to 6.47 ± 1.87 µmol O_2_/g·h) and began to decrease at 60 % air sat. (164 µmol O_2_/L; 5.73 ± 1.38 µmol O_2_/g·h). Beginning at 40 % air sat. (109 µmol O_2_/L), MMR followed an oxygen conforming pattern. *Ṁ*O_2_ measured at 30, 25 and 20% O_2_ were significantly lower than at treatments 50% to 100% air sat.. MMR matched SMR at 21.6 % air sat. (59 µmol O_2_/L, 4.5 kPa). During the acclimation period to 25 – 20% air sat., four of the seven fish lost equilibrium and one fish died at the end of the acclimation period.

#### 3.2.4 *P*_crit_, P_crit-max_ and LOL curve

At 2.4 °C, *B. saida’s P*_crit_ accounted for 5.9 % air saturation (21.17 µmol O_2_/L) (Figures 3-5, Table 1). Only the intersection point of the MMR and SMR regression lines could be used for *P*_crit_ calculations, since the regression line calculated for the RMR had no intersection point with the SMR regression line. *P*_crit-max_ amounted to 56.0 ± 2.8 % ambient oxygen saturation (11.8 ± 0.6 kPa O_2_), resulting in Δ*P*_crit-max_ – *P*_crit_ of 50.1 % air sat (10.6 kPa)

**Figure 3.**
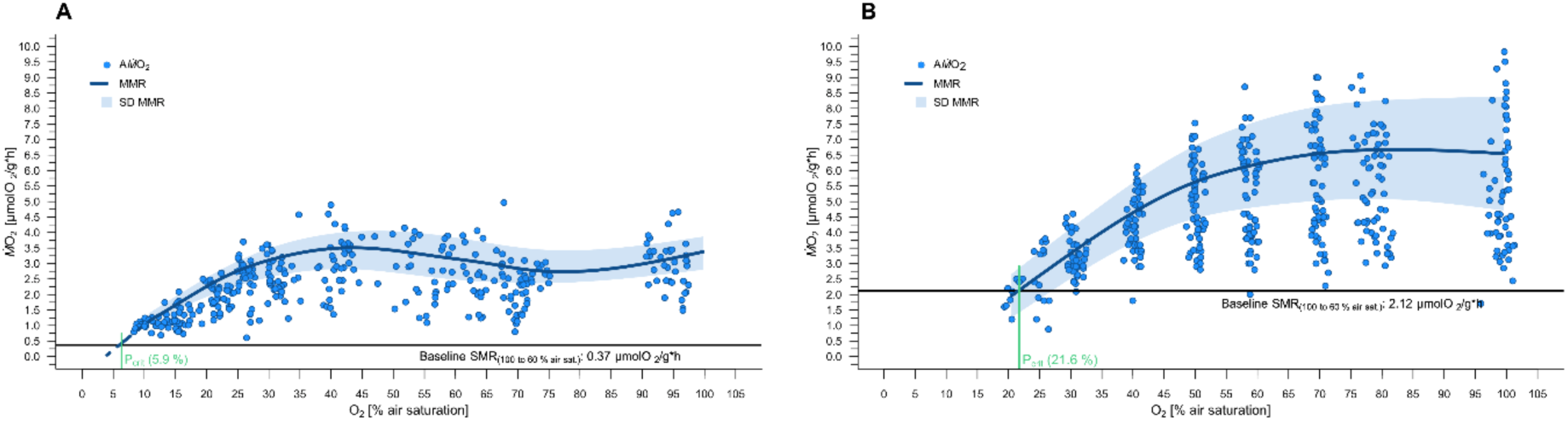
Active metabolism and maximum metabolic rate with progressing hypoxia at 2.4 and 10.0 °C. The oxygen consumption ((*Ṁ*O_2_) in µmol O_2_/g·h) over oxygen saturation (in % air saturation) is displayed as active oxygen consumption (blue circles, A*Ṁ*O_2_) and maximum metabolic rate per treatment group (dark blue line, MMR as polynominal regression) with standard deviation (blue shaded area, SD MMR). At the lower end of the oxygen range the critical oxygen saturation (*P*_crit_, in % air sat.) is included (green line). The experiments were terminated either by reaching *P*_crit_ or by an increasing number of fish refusing to swim, ending the swimming trial. As in Figure 1, the baseline SMR included as black line (Baseline SMR). **A:** AMR at 2.4 °C. 361 single *Ṁ*O_2_ data points representing 60 experimental runs of 30 individuals. A *P*_crit_ of 5.9 % air sat. (21.2 µmol O_2_ /L) was calculated. **B:** AMR at 10.0 °C. 431 single *Ṁ*O_2_ data points representing 58 experimental runs of 23 individuals. A *P*_crit_ of 21.6 % air sat. (59.0 µmol O_2_ /L) was calculated.

**Figure 4.**
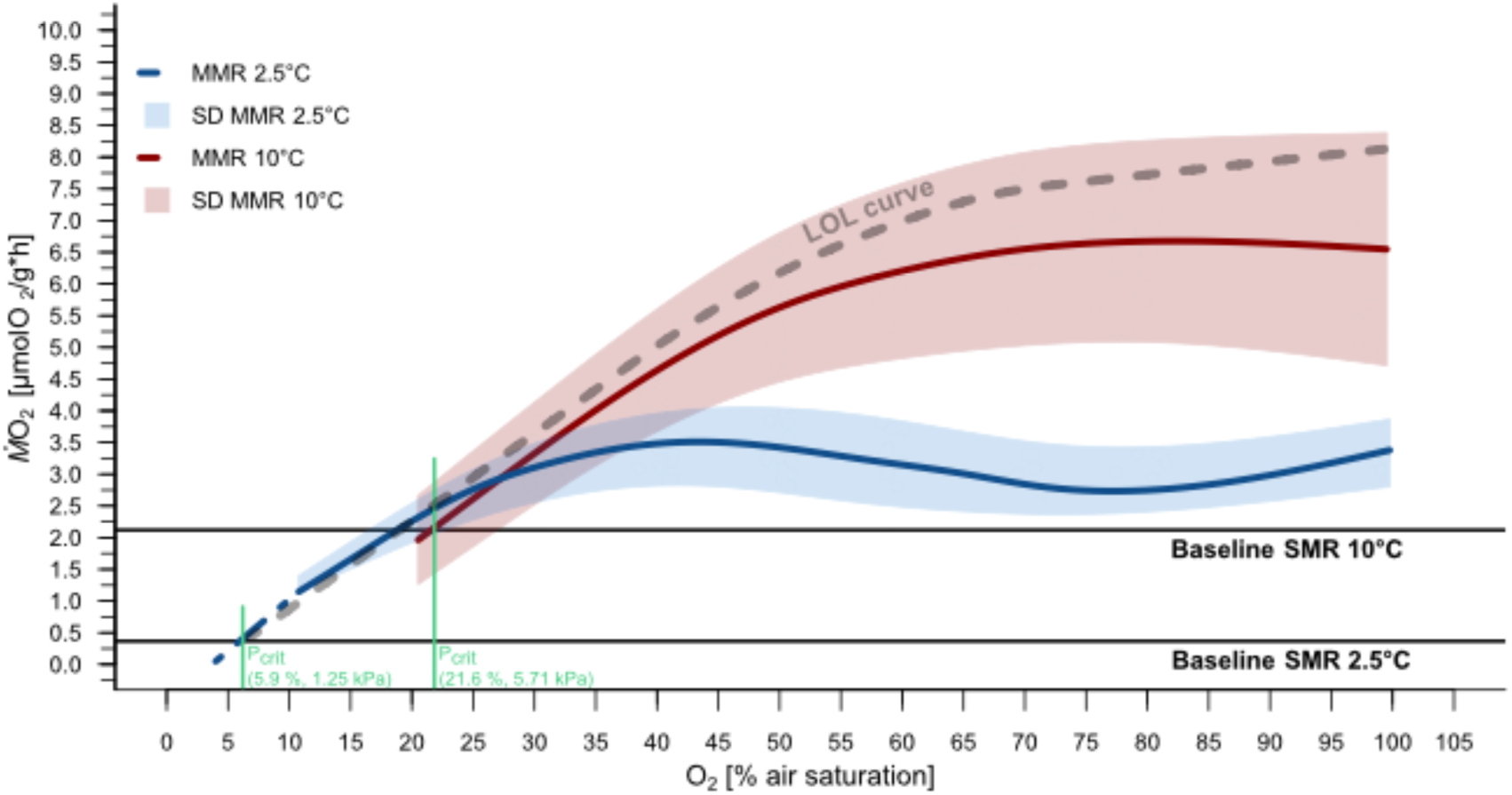
Limiting oxygen level (LOL) curve after Claireaux and Chabot (2016). The LOL curve (grey dashed line), a continuum of limiting oxygenation levels is prognosticated from the maximum metabolic rates (MMR, (*Ṁ*O_2_) in µmol O_2_/g·h) at 2.4 °C (blue line, with a polynomial regression of the *Ṁ*O_2_ recordings from 10 – 25 % air sat. for *P*_crit_ calculation: dashed blue line) and 10.0 °C (red line) with their MMRs corresponding standard deviation (blue and red shaded areas), *P*_crit_ (light green circles; 5.9 % air sat. at 2.4 °C and 21.6 % air sat. at 10.0 °C) and baseline SMRs (black lines; 0.37 µmol O_2_/g·h at 2.4 °C and 2.12 µmol O_2_/g·h at 10.0 °C).

**Figure 5.**
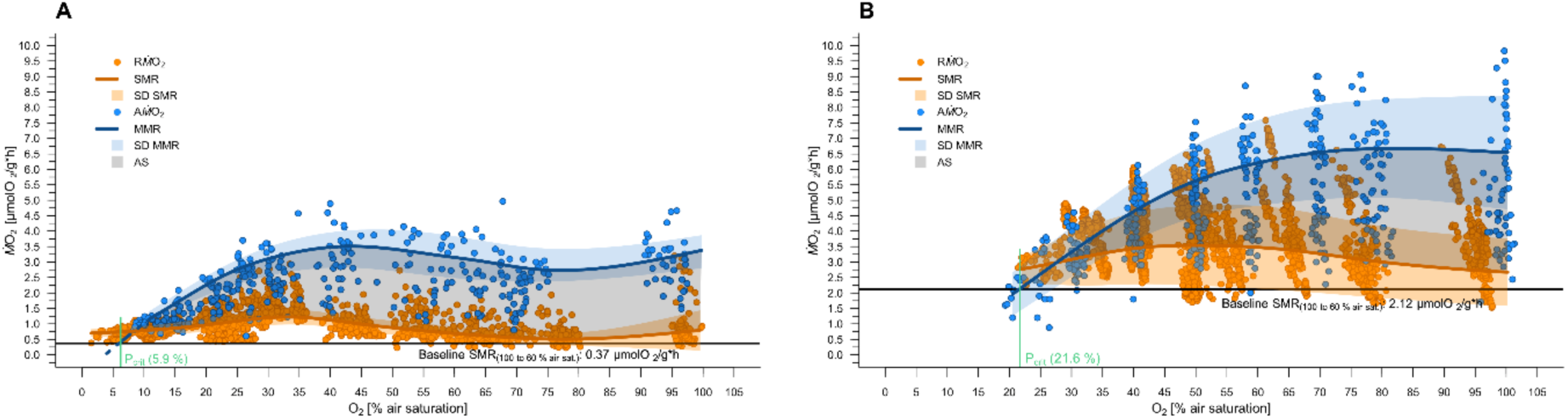
Routine-, and active metabolism, standard-, and maximum metabolic rate and corresponding aerobic scope with progressing hypoxia at 2.4 and 10.0 °C. The oxygen consumption ((*Ṁ*O_2_) in µmol O_2_/g·h) over oxygen saturation (in % air saturation) is displayed as active oxygen consumption. This figure combines Figure 1 and Figure 2 and supplements them by the aerobic scope (grey shaded area, AS), the difference between MMR and SMR for both temperature treatments. **A:** Measurements of routine and active *Ṁ*O_2_ and corresponding AS at 2.4 °C. **B:** Measurements of routine and active *Ṁ*O_2_ and corresponding AS at 10.0 °C.

In the warm acclimated group, an intersection point between MMR and baseline SMR was visible, resulting in a *P*_crit_ of 21.6 % air sat. (4.5 kPa O_2_).

*P*_crit-max_ amounted to 68 ± 3.4 ambient oxygen saturation (14.2 ± 0.7 kPa O_2_), resulting in Δ*P*_crit-max_ – *P*_crit_ of 46.4 % air sat (9.7 kPa).

#### 3.2.4 Absolute and factorial aerobic scope

In the cold, the aerobic scope in figure 5A significantly decreased with decreasing oxygen saturation (p < 0.001). At 10.0 °C (Figure 5B), AS displayed a similar pattern and also decreased significantly with decreasing oxygen concentration (p < 0.05).

At both temperatures, a stepwise decrease of AS was observed. At 2.4 °C, AS accounted for 2.49 ± 0.77 µmol O2/g·h within an oxygen range of 95.53 ± 4.52 % air sat. (333.59 ± 15.07 µmol O2/L) to 43.65 ± 2.65 % air sat. (153.86 ± 4.08 µmol O2/L). In comparison the AS decreased in the second group (from 33.06 ± 2.11 to 21.15 ± 1.46 % air sat. (115.00 ± 2.42 to 73.57 ± 1.07 µmol O2/L), p < 0.001), AS decreased to 1.56 ± 0.50 µmol O2/g·h. In the lowest *P*O_2_ group (from 13.95 ± 1.41% to 10.96 ± 1.03 % air sat. (48.51 ± 0.68 to 38.36 ± 0.39 µmol O2/L), p < 0.001), also the lowest AS of 0.55± 0.24 µmol O2/g·h was calculated. The highest value for aerobic scope, 2.77 ± 0.48 µmol O2/g·h, was recorded at 43.65 ± 2.65 % air saturation.

At 10.0 °C, AS remained stable between 100-70% O_2_ saturation with a mean of 3.58 ± 1.75 µmol O_2_/g·h. AS at 60 and 50% air sat. was significantly decreased compared to 80% air sat. (p=0.001, p=0.019) with an AS of 2.53 ± 1.47 µmol O_2_/g·h and 2.76 ± 1.53 µmol O_2_/g·h at 60 and 50% air sat., respectively. AS did not differ significantly between the three lowest *P*O_2_ levels tested (p=0.46) with a mean AS of 1.34 ± 0.94 µmol O_2_/g·h.

In line with absolute AS, factorial AS was also significantly negatively affected by decreasing oxygen concentrations (Figure 6).

**Figure 6.**
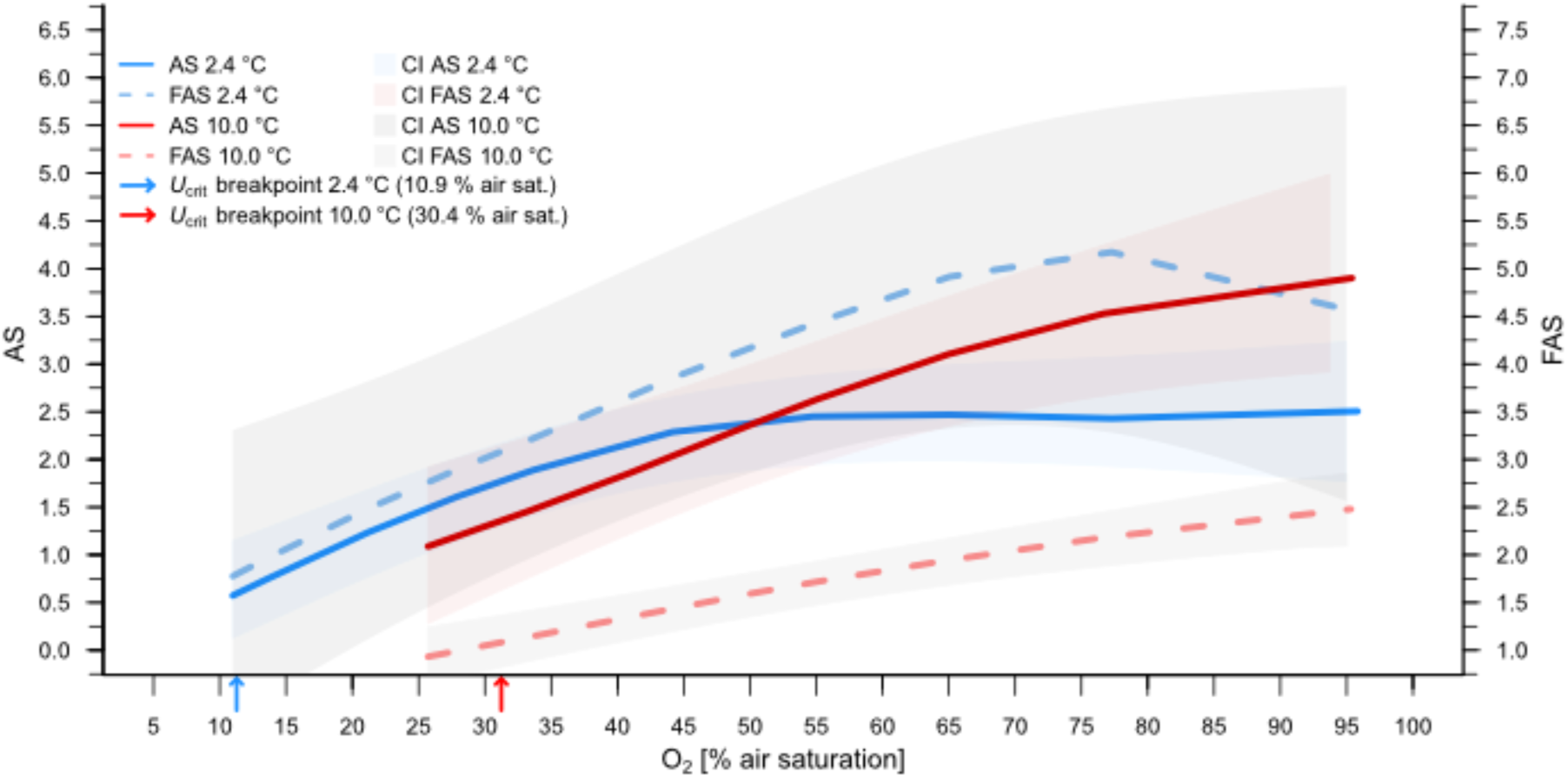
Aerobic scope (AS) and factorial aerobic scope (FAS). The AS for 2.4 °C (solid blue line) and 10.0 °C (solid red line) and FAS for respectively 2.4 °C (dashed blue line) and 10.0 °C (dashed red line) with their confidence intervals (CI) are shown. The blue and red arrows on the x-axis indicate the breakpoint where fish had to reduce their maximum swimming speed (U_crit_).

The factorial aerobic scope (MMR/SMR; FAS) sank to <50% in the warm acclimated fish compared to that at 2.4 °C (Figure 6). This follows that warm AS was initially larger at 100 % air sat. but also started to decrease earlier (70 % vs. 50 %), meeting with cold AS around 50 % air sat and running in parallel along an oxyconforming line until finally reaching *P*_crit_ at 21.6 % air sat.

Interestingly, U_crit_ remained constant over a far wider *P*O_2_ range than AS could be maintained, visibly decreasing only at 10% and 30% air sat. in the cold and warm treatments, respectively. This corresponds to about 20 % of total AS in the cold and 35% in the warm.

### 3.3 Swimming performance

In general, we observed a decline of swimming activity with progressive hypoxia at both temperatures.

At 2.4 °C, the total active swimming time significantly decreased with decreasing oxygen saturation (p = 0.017). Although we were not able to detect such a significant effect for the warm acclimated fish (p=0.72) we found the lowest swimming activity at 25% air sat.. Furthermore, two fish lost equilibrium during the acclimation period to this *P*O_2_ treatment. At 20% air sat., another two fish lost equilibrium during the acclimation period and one died.

#### 3.3.1 MR at different water velocities

*Ṁ*O_2_ increased significantly with increasing water velocities for both temperature treatments (p < 0.001).

The highest *Ṁ*O_2_ at 2.4 °C was detected at 2.60 BL/sec with an average of 4.94 ± 0.80 µmol O_2_/g·h. With increasing water velocities, an increasing number of fish refused to swim (point of U_crit_). The lower sample size (n=59) at 1.40 and 1.55 BL/sec was due to measurement disturbances of the oxygen optodes so that these measurements had to be excluded from further analysis.

At 10.0 °C, metabolic rates increased significantly with water velocity up to 2.9 BL/sec and a mean *Ṁ*O_2_ of 6.47 ± 1.49 µmol O_2_/g·h (Table 1). With increasing water velocities, an increasing number of fish refused to swim, resulting in only three individuals reaching the maximum swimming speed of 3.5 BL/sec.

#### 3.3.2 Gait transition speed **U_gait_**

At 2.4 °C, *P*O_2_ had no significant influence on U_gait_ (p= 0.86).

Between 100 and 30% air saturation (357.57 – 112.07 µmol O_2_/L) all fish displayed burst swim behaviour and a rather constant U_gait_ of 2.25 BL/sec. Below 30 % air saturation, the total occurrence of burst swim behaviour was reduced with progressive hypoxia (Table 1). In the 20 % treatment (21.70 ±0.57 % air saturation, 77.75 ± 0.44 µmol O_2_/L), only three individuals displayed burst swim behaviour at all. The remaining three Polar cod stopped swimming without detectable burst actions. In the 10 % air sat. treatment (10.67 % air saturation; 39.12 ± 0.59 µmol O_2_/L and U_gait_=1.87 BL/sec), only one Polar cod out of six displayed burst swim behaviour. Additionally, the burst swimming activity, determined as the total number of bursts (p = 0.025) and the number of bursts per minute (p = 0.054) decreased significantly with decreasing *P*O_2_ (Table 1). The maximum number of 14.56 ± 9.33 bursts per minute was achieved in the 30 % air sat. treatment (Table 1).

Although statistical analysis revealed a global significant effect of *P*O_2_ on U_gait_ at 10.0 °C (Kruskal-Wallis, p=0.02), this was not confirmed by the post-hoc test (Wilcoxon-Test) between the individual oxygen treatments. Generally, U_gait_ was relatively stable over a *P*O_2_ range of 100 to 60% air sat. (274 – 164 µmol O_2_/L; median: max: 2.1 BL/sec, min: 1.92 BL/sec) with a maximum of 2.05 ± 0.17 BL/sec at 60 % air sat. (Table 1), yet significantly decreased compared to 2.4 °C (p < 0.001). The number of animals which showed burst swimming behaviour varied between the *P*O_2_ treatments (100-30 % air sat: n=5-6) and decreased to an n=2 at 25 % air sat.. At 20 % air sat, no more burst swimming behaviour occurred.

The burst swimming activity (total number of bursts) and the number of bursts per minute did not reveal a significant *P*O_2_ effect (p = 0.23, and p = 0.42). Nonetheless, despite their variability, both parameters showed highest values at 40 % air sat. with 226.33 ± 316.60 total number of bursts and corresponding 5.86 ± 5.18 bursts per minute, followed by a decrease at 30 % air sat.

#### 3.3.3 Critical swimming speed (**U_crit_**)

At 2.4 °C, U_crit_ was maintained throughout all treatments, only the U_crit_ values in the 10% air sat. treatment were significantly lower than under normoxia (p=0.031). We found a similar picture at 10.0 °C, with U_crit_ remaining relatively stable over an O_2_ range from 100 to 40% air sat. (max: 2.61 BL/sec, min: 2.50 BL/sec) but significantly lower than at 2.4 °C (p < 0.001).

## 4 Discussion

There are very few studies that have determined the critical oxygen saturation (*P*_crit_) and examined the influence of ambient oxygen level upon aerobic scope (AS) of polar fish [e.g. 42, 61]. To close this gap in knowledge, we measured metabolic rates in Polar cod (*B. saida*) under progressive hypoxia at 2.4 °C and after warm acclimation to close to their thermal limit (10.0 °C). We observed very clear and stable patterns that were similar in both thermal regimes. Over the whole range of their SMR and in part also for MMR (above 40 % and 70 % air saturation, respectively), Polar cod displayed oxygen regulating behaviour under progressive hypoxia, with SMR never below aerobic baseline metabolism. As a result, we found *P*_crit_ to be unexpectedly low at 5.9 % air saturation (21.2 µmol O_2_/L, 1.3 kPa) and AS to be maintained in hypoxia until about 45% air saturation (156.9 µmol O_2_/L) at typical habitat temperatures. Closer to critical temperatures, aerobic scope could only be maintained down to about 70 % air saturation (191.5 µmol O_2_/L) and *P*_crit_ rose to 21.6 % air saturation (59 µmol O_2_/L).

In comparison, literature values for Greenland cod (*Gadus ogac*) and Atlantic cod (*Gadus morhua*) show temperature and life stage dependent *P*_crit_ ranging from 31.7 % to 41.6 % air saturation at 1 °C for adult Greenland cod, and 16.5-28.5 % (at 5°C) to 15.5 % (at 10°C) for juvenile Atlantic cod [17, 42, 60, 62, 63]. This reflects the different metabolic thermal optima observed for growth and exercise in Polar and Atlantic cod [22].

The tolerance of Polar cod to these low oxygen saturations is certainly supported by their naturally very low SMR (baseline SMR at 2.4 °C = 0.37 µmol O_2_/g·h), indicating that these fish do not need much oxygen to sustain basic metabolic functions in their cold and thermally stable habitat. This may have set the evolutionary basis for hypoxia tolerance as similar energy conservation mechanisms can be used under both hypoxia and hypothermia [64-66].

It is not all about tolerating, though: in the cold, we observed SMR to be actively up-regulated between 45 and 35 % air saturation and MMR to decline steadily, which progressively narrowed AS. Only below 35 % air saturation, SMR and MMR followed an oxyconforming pattern with decreasing *P*O_2_ before merging at *P*_crit_. Typically, *Ṁ*O_2_ would decline below baseline SMR at *P*_crit_ when fish enter their anaerobic “scope for survival” [52]. However, *Ṁ*O_2_ always remained above baseline SMR in Polar cod before they lost equilibrium at 2 % air saturation. This resulted in the need to calculate a theoretical *P*_crit_ [according to 52] as an intersection of MMR with baseline SMR was not observed. We therefore conclude that Polar cod have a very limited anaerobic capacity. In line with this assumption is the decrease of anaerobically fuelled burst swim behaviour with decreasing oxygenation (c.f. ‘burst swimming’, Table 1), along with a decrease in active swimming duration. Kunz et al. [23] already noted that Polar cod showed only rare and very brief periods of burst swim activity under different temperature/hypercapnia treatments and determined an anaerobically fuelled swimming capacity of only 0−5.7 %. Earlier studies have reported a generally low anaerobic capacity for polar fish [67, 68]. Our present data point in the same direction and corroborate the notion that in Polar cod these short bouts of anaerobic burst swimming are only possible under conditions where almost instantaneous aerobic recovery of white swimming musculature is possible [cf. 69].

Thus, there are clear indications for a maximisation of aerobic metabolism in Polar cod by active upregulation of SMR under hypoxic conditions. When compared to other fish, such as European sea bass (*Dicentrarchus labrax*), other gadoids (e.g. *Gadus morhua, Gadus ogac*), common sole (*Solea solea)* and turbot (*Psetta maxima*) [reviewed by 70], which all display declining SMR and AS with decreasing oxygen content, the regulatory capacity to maintain AS to such low *P*O_2_ levels therefore appears extraordinary in Polar cod. For example, SMR of arctic Greenland cod (*Gadus ogac*) starts to oxyconform around 45% air sat. [70 mmHg 42], falling below baseline SMR, while Polar cod upregulate their *Ṁ*O_2_ in this range and never fall below baseline SMR. This upregulation is further supported by an exceptionally low baseline SMR, which is among the lowest *Ṁ*O_2_ that have been measured in fish, pointing at relatively little energy required for oxygen extraction and perfusion in oxygen rich polar waters – which may be supported by a certain degree of cutaneous respiration that is frequently found in polar fish [e.g. 71, 72, 73]. Increased O_2_ extraction costs under progressive hypoxia are evidenced by rising metabolic rates below 50 % air sat. This intermediate increase in baseline *Ṁ*O_2_ appears as a species-specific strategy to deal with decreasing oxygen levels and results in one of the lowest *P*_crit_ we are aware of for fish at their respective habitat temperature [Pcrit 1.25 kPa, cf. 60]. Consequently, MMR (and thus aerobic scope) is stabilized down to relatively low *P*O_2_, which may be of even greater importance for Polar cod in everyday life.

At first sight, the presence of a strongly developed hypoxia tolerance in a polar fish may appear counterintuitive in light of the general paradigm that hypoxic water layers are mostly associated with temperate and tropical waters. Nonetheless, Polar cod can experience hypoxic conditions in the Svalbard fjord systems and may even be evolutionarily adapted to them: as pointed out above, high-silled semi-closed fjords depend on winter circulation driven by ice formation for bottom water exchange and re-oxygenation, leading to local hypoxia in these deep cold fjord basins where Polar cod spend the summer months [10, 12]. Furthermore, Polar cod is known to occur in very large schools of up to 900 million individuals [20, 74, 75] and generally schooling leads to progressive hypoxia towards the center of the schools [76-79]. Polar cod have thus been subjected to an evolutionary need to develop maximised oxygen extraction capacities to reduce ventilatory and general metabolic costs, which is also reflected in a low *P*_crit_.

However, the pertinent question remains whether the observed responses can be summarized under classic hypoxia tolerance, as we a) did not observe any metabolic downregulation and b) no anaerobic component of the hypoxia response in Polar cod, which are usually put forward in the definition of hypoxia tolerance [64, 66, 80]. Therefore, we would tend to describe the observed metabolic response to hypoxia rather as metabolic hypoxia compensation than hypoxia tolerance as the mechanisms involved here actively seek to improve oxygen supply instead of (anaerobically) tolerating hypoxia through metabolic depression.

Evolution of this metabolic compensation strategy was possible in the generally cold and oxygen rich Arctic waters, but also necessary, as the Arctic regions around Svalbard underwent several climate warming phases in their past, with winter mean air temperatures being up to 10 K warmer than nowadays [81], with ice free winters and hypoxic fjord bottom waters as a possible consequence.

In light of the ongoing climate change, the question arises whether Polar cod can compensate for warming and hypoxic conditions stronger than the warm phases of the past?

Over the past 100 years, Isfjorden (the connecting fjord between Billefjorden and the Ocean) has warmed by 1.9 K [82], Arctic surface air temperatures are projected to rise by 5-10 K [6, 83] and Arctic sea surface temperatures (SST) are projected to rise by up to 0.5 K per decade [84].

We therefore conducted a second set of identical hypoxia experiments at 10.0 °C with a group of fish that had been long-term acclimated to this temperature in order to see if and how far *P*_crit_ would shift and whether the underlying mechanisms were still present.

In general, we observed similar patterns in SMR and MMR after warm acclimation compared to 2.4 °C, albeit more compressed. Below T_crit_, Q10 would predict a doubling of baseline metabolic rate between 2 and 10.0 °C, if metabolic compensation during acclimation is disregarded. In the present experiment, the 10.0 °C acclimated fish displayed a 5-fold higher baseline *Ṁ*O_2_ compared to 2.4 °C, indicating that there was no metabolic compensation during acclimation and instead metabolic costs had risen exponentially.

These high costs can be due to metabolic/mitochondrial inefficiency above Polar cod’s metabolic and growth optima on the one hand [22, 25] and rising costs of the circulatory system on the other [cf. 85]. We found MMR to be increased two-fold in the warm compared to 2.4 °C, which can be an acute effect of temperature or a result of acclimation to 10.0 °C (figure 5, table 1). In comparison to the observed 5-fold rise in SMR, this indicates a physical limitation of either oxygen uptake and cardio-vascular distribution or ATP turnover and demand. As active swimming time is not reduced in the warm compared to 2.4 °C, we can probably rule out oxygen distribution and ATP supply, and thus have to attribute the limited MMR capacity in the warm to limitations of the aerobic swimming musculature.

Further, there appears to be a common ‘oxyconforming line’ across temperatures, to which all MMR curves adhere at their lower limits, supporting the concept of limiting oxygen levels [Figure 4, cf. 52, 86].

The higher initial MMR in the warm acclimated fish also led to an increase of absolute AS in the range between 100 and 50 % air sat. (cf. Table 1, Figure 6). However, comparing absolute AS at different temperatures can be unrealistic, as it ignores rising baseline metabolic costs that claim a higher percentage of metabolic scope [87]. These reflect generally higher maintenance costs in the warm. Therefore, factorial aerobic scope (MMR/SMR; FAS) sinks to < 50 % in the warm acclimated fish compared to that at 2.4 °C, despite the increased absolute AS (Figure 6). This follows that warm AS is initially larger at 100 % air sat. but also starts to decrease earlier (70 % vs. 50 %), meeting with cold AS around 50 % air sat. (Figure 6) and running in parallel along an oxyconforming line until finally reaching *P*_crit_ at 21.6 % air sat.

Nonetheless, as a result of the increased MMR *P*_crit-max_ could be stabilized in the same range at both temperatures (2.4 °C: 11.8 ± 0.6 kPa and 10.0 °C: 14.2 ± 0.7 kPa) and the oxyconforming range between *P*_crit_ and *P*_crit-max_ was virtually constant (10.6 vs. 9.7 kPa). This points at similar oxyconforming processes being employed in both groups, with WA fish reaching their limits a little earlier due to higher SMR and lower relative oxygen saturation in the water.

A *P*_crit_ of 21.6 % air sat. (59.1 µmol O_2_/L) at 10.0 °C is surprisingly similar to the *P*_crit_ of its larger relative Atlantic cod, *Gadus morhua* at this temperature, 23% air sat. (62.9 µmol O_2_/L) [62, 70], which is invading the warming Arctic waters in increasing numbers [17, 88, 89]. Thus in a warmer future, Polar cod would lose their advantage of a substantially lower *P*_crit_ over invading species and predators. This is also mirrored in behavioural changes at 10.0 °C: despite lower FAS, active swimming time was constantly high throughout the oxygen range as opposed to a reduction in active swimming time with decreasing oxygen levels at 2.4 °C.

In our experiments we tried to access *B. saida’s* anaerobic capacity in detecting behavioural changes under swimming exercise paired with progressive hypoxia. It became clear that *B. saida* has a generally low capacity for anaerobic swimming, whereas the overall swimming capacity is comparable to its larger relative and invading predator, *Gadus morhua*, which reaches similar maximum swimming speeds at 5 °C [cf. 90, 91, 92]. Interestingly, U_crit_ remained constant over a far longer *P*O_2_ range than AS could be maintained, visibly decreasing only at 10 % and 30 % air sat. in the cold and warm treatments, respectively. This corresponds to about 20 % of total AS in the cold and 35 % in the warm. As such, swimming is compromised only very late when aerobic scope is reduced during progressive hypoxia, indicating that it has a high priority in energy allocation in Polar cod. This indirectly also indicates that not more than 20-35 % of aerobic scope is fuelled into swimming, which translates into a relatively modest cost of swimming at these temperatures [93].

## 5 Conclusion

In conclusion, we observed that Polar cod can maintain their AS over a wide oxygen range even under warming. This appears extraordinary compared to other Gadoids [reviewed by 70]. We did not detect any metabolic depression or anaerobic component of Polar cod’s hypoxia response at both temperatures, which was mirrored in a constant maximum swimming and gait transition speed even at oxygen saturations close to its *P*_crit_. The fact that these locomotory traits are maintained even under decreasing AS indicates a clear priority of swimming in energy allocation.

The absence of anaerobic capacities suggests that Polar cod adopted a strategy to prevent systemic hypoxia, which can also be found in other obligate aerobic fish species, such as salmonids (pers. obs. G. Claireaux, D. Chabot, C. Audet). However, in contrast to salmonids, Polar cod have a much lower *P*_crit_ (0 °C: 5.9 % air sat. (21.2 µmol O_2_/L, 1.3 kPa); 10.0 °C: 21.6 % air sat. (59.1 µmol O_2_/L, 4.5 kPa)) and baseline SMR (2.4 °C: 0.37 µmol O_2_/g·h, 10.0 °C: 2.12 µmol O_2_/g·h), implying that these fish generally do not need much oxygen to sustain basic metabolic functions [also to deal with periods of winter starvation, see 94]. Metabolic hypoxia compensation as displayed by Polar cod is a helpful strategy in the short run, though – in the long run, the ‘compensation’ aspect means that there is a trade-off and swimming priority comes at a long-term metabolic cost, e.g. fecundity or reproduction as such. A reduced fecundity (i.e., number of eggs in females), maturity at an earlier age and smaller body size has already been reported for Polar cod originating from the warmer Atlantic sector compared to those from more Arctic water influenced regions [95]. Together with a potential mismatch between larval development and prey availability caused by delayed sea ice formation [24, 96], these *P*O_2_-dependent trade-offs may strengthen a negative influence on Polar cod reproduction, population size and abundance.

Our study thus indicates that Polar cod rather compensate metabolic hypoxia than employ classic hypoxia tolerance strategies. Their ability to maintain metabolic functions under extreme hypoxic conditions will help Polar cod in a warmer future with a progressive loss of sea ice formation in winter and consequent decrease of oxygen content [97] to spend their summers in less oxygenated or even hypoxic cold water basins in the fjords interior. This provides a potential shelter from less hypoxia resistant predacious fish, which are likely to remain in the surface layers [see 42, 62, 70, 98]. However, these low-oxygen basins in the fjords can only serve as protective zones for Polar cod for the next decade(s) as long as summer water temperatures still stay cold – despite relatively constant *P*_crit-max_, their *P*_crit_ will increase at warmer temperatures and this physiological advantage of Polar cod over invading species and predators will be lost.

## 7 Appendix

## 8 Acknowledgements

This project was funded through the AWI research program POF IV Topic 06 subtopic 02 and is a contribution to the EU project ACTNOW (HORIZON-CL6-2021-BIODIV-01 project no. 101060072). This project would not have been possible without the continuous scientific support, fruitful discussions and mentoring of Prof. Dr. Guy Claireaux, to whom we are grateful and indebted. We would like to thank the crews of RV Heincke (AWI, funding No. AWI_HE519_01 and AWI_HE560_01) for their support in animal collection. Further, we would like to thank Fredy Veliz Moraleda and Amirhossein Karamyar for technical assistance with the experimental set-up and animal welfare.

## 9 Ethical approval

All procedures performed in the present study were in accordance with the ethical standards of the federal state of Bremen, Germany, and were approved under the reference number *Tierversuch Nr. 160 500-427-103-7/2018-1-5*.

## Notes

### Competing Interest Statement

The authors have declared no competing interest.

## 6 References

1. Serreze MC, Barry RG: **Processes and impacts of Arctic amplification: A research synthesis**. Global and Planetary Change 2011, 77(1-2):85–96.

2. Winton M: **Does the Arctic sea ice have a tipping point?** Geophysical Research Letters 2006, 33(23).

3. Kwok R, Untersteiner N: **The thinning of Arctic sea ice**. Phys Today 2011, 64(4):36–41.

4. Overland JE, Wang M: **When will the summer Arctic be nearly sea ice free?** Geophysical Research Letters 2013, 40(10):2097–2101.

5. IPCC: Climate Change 2014: **Synthesis Report. Contribution of Working Groups I, II and III to the Fifth Assessment Report of the Intergovernmental Panel on Climate Change. With assistance of Core Writing Team, R. K. Pachauri and L. A. Meyer**. Geneva, Switzerland. 2014.

6. IPCC: IPCC Special Report on the Ocean and Cryosphere in a Changing Climate [H.-O. Pörtner, D.C. Roberts, V. Masson-Delmotte, P. Zhai, M. Tignor, E. Poloczanska, K. Mintenbeck, A. Alegría, M. Nicolai, A. Okem, J. Petzold, B. Rama, N.M. Weyer (eds.)]. Cambridge University Press, Cambridge, UK and New York, NY, USA 2019:755 pp.

7. IPCC: Climate Change 2022: **Impacts, Adaptation, and Vulnerability. Contribution of Working Group II to the Sixth Assessment Report of the Intergovernmental Panel on Climate Change**. Cambridge, UK and New York, NY, USA: Cambridge University Press; 2022.

8. Collins M, Knutti R, Arblaster J, Dufresne J-L, Fichefet T, Friedlingstein P, Gao X, Gutowski WJ, Johns T, Krinner G: **Long-term climate change: projections, commitments and irreversibility**. In: Climate Change 2013-The Physical Science Basis: Contribution of Working Group I to the Fifth Assessment Report of the Intergovernmental Panel on Climate Change. Cambridge University Press; 2013: 1029–1136.

9. Nilsen F, Cottier F, Skogseth R, Mattsson S: **Fjord–shelf exchanges controlled by ice and brine production: The interannual variation of Atlantic Water in Isfjorden, Svalbard**. Continental Shelf Research 2008, 28(14):1838–1853.

10. Szczuciński W, Zajączkowski M, Scholten J**: Sediment accumulation rates in subpolar fjords–Impact of post-Little Ice Age glaciers retreat, Billefjorden, Svalbard**. Estuarine, Coastal and Shelf Science 2009, 85(3):345–356.

11. Renaud PE, Wlodarska-Kowalczuk M, Trannum H, Holte B, Weslawski JM, Cochrane S, Dahle S, Gulliksen B**: Multidecadal stability of benthic community structure in a high-Arctic glacial fjord (van Mijenfjord, Spitsbergen)**. Polar Biol 2007, 30(3):295–305.

12. Promińska A, Falck E, Walczowski W: **Interannual variability in hydrography and water mass distribution in Hornsund, an Arctic fjord in Svalbard**. Polar Research 2018, 37(1):1495546.

13. Cottier F, Nilsen F, Skogseth R, Tverberg V, Skarðhamar J, Svendsen H**: Arctic fjords: a review of the oceanographic environment and dominant physical processes**. Geological Society, London, Special Publications 2010, 344(1):35–50.

14. Cottier F, Tverberg V, Inall M, Svendsen H, Nilsen F, Griffiths C: **Water mass modification in an Arctic fjord through cross-shelf exchange: The seasonal hydrography of Kongsfjorden, Svalbard**. Journal of Geophysical Research: Oceans 2005, 110(C12).

15. Matear R, Hirst A: **Long-term changes in dissolved oxygen concentrations in the ocean caused by protracted global warming**. Global Biogeochemical Cycles 2003, 17(4).

16. Keeling RF, Kortzinger A, Gruber N: **Ocean Deoxygenation in a Warming World**. Annu Rev Mar Sci 2010, 2:199–229.

17. Renaud PE, Berge J, Varpe Ø, Lønne OJ, Nahrgang J, Ottesen C, Hallanger I: **Is the poleward expansion by Atlantic cod and haddock threatening native polar cod, Boreogadus saida?** Polar Biology 2012, 35(3):401–412.

18. Eriksen E, Ingvaldsen RB, Nedreaas K, Prozorkevich D: **The effect of recent warming on polar cod and beaked redfish juveniles in the Barents Sea**. Regional Studies in Marine Science 2015, 2:105–112.

19. Majewski AR, Walkusz W, Lynn BR, Atchison S, Eert J, **Reist JD: Distribution and diet of demersal Arctic Cod, *Boreogadus saida*, in relation to habitat characteristics in the Canadian Beaufort Sea**. Polar Biology 2016, 39(6):1087–1098.

20. Crawford R, **Jorgenson J: Quantitative studies of Arctic cod (*Boreogadus saida*) schools: important energy stores in the Arctic food web**. Arctic 1996:181–193.

21. Crawford RE, Vagle S, Carmack EC: **Water mass and bathymetric characteristics of polar cod habitat along the continental shelf and slope of the Beaufort and Chukchi seas**. Polar Biology 2012, 35(2):179–190.

22. Kunz KL, Frickenhaus S, Hardenberg S, Johansen T, Leo E, Pörtner H-O, Schmidt M, Windisch HS, Knust R, **Mark FC: New encounters in Arctic waters: a comparison of metabolism and performance of polar cod (*Boreogadus saida*) and Atlantic cod (*Gadus morhua*) under ocean acidification and warming**. Polar Biology 2016, 39(6):1137–1153.

23. Kunz KL, Claireaux G, Pörtner H-O, Knust R, **Mark FC: Aerobic capacities and swimming performance of polar cod (*Boreogadus saida*) under ocean acidification and warming conditions**. Journal of Experimental Biology 2018, 221(21):jeb184473.

24. Leo E, Dahlke FT, Storch D, Portner HO, Mark FC: **Impact of Ocean Acidification and Warming on the bioenergetics of developing eggs of Atlantic herring *Clupea harengus***. Conservation Physiology 2018, 6.

25. Leo E, Kunz KL, Schmidt M, Storch D, Portner HO, **Mark FC: Mitochondrial acclimation potential to ocean acidification and warming of Polar cod (*Boreogadus saida*) and Atlantic cod (*Gadus morhua*)**. Front Zool 2017, 14:21.

26. Schmidt M, Gerlach G, Leo E, Kunz KL, Swoboda S, Pörtner H-O, Bock C, **Storch D: Impact of ocean warming and acidification on the behaviour of two co-occurring gadid species, *Boreogadus saida* and *Gadus morhua*, from Svalbard**. Marine Ecology Progress Series 2017, 571:183–191.

27. Drost H, Carmack E, **Farrell A: Upper thermal limits of cardiac function for Arctic cod *Boreogadus saida*, a key food web fish species in the Arctic Ocean**. Journal of fish biology 2014, 84(6):1781–1792.

28. Drost H, Fisher J, Randall F, Kent D, Carmack E, Farrell A: **Upper thermal limits of the hearts of Arctic cod Boreogadus saida: adults compared with larvae**. Journal of fish biology 2016, 88(2):718–726.

29. Griffith GP, Hop H, Vihtakari M, Wold A, Kalhagen K, Gabrielsen GW: **Ecological resilience of Arctic marine food webs to climate change**. Nature Climate Change 2019, 9(11):868–872.

30. Sampaio E, Santos C, Rosa IC, Ferreira V, Portner HO, Duarte CM, Levin LA, Rosa R: **Impacts of hypoxic events surpass those of future ocean warming and acidification**. Nat Ecol Evol 2021, 5(3).

31. Limburg KE, Breitburg D, Levin LA: **Ocean deoxygenation - a climate-related problem**. Front Ecol Environ 2017, 15(9):479–479.

32. Limburg KE, Breitburg D, Swaney DP, Jacinto G**: Ocean Deoxygenation: A Primer**. One Earth 2020, 2(1):24–29.

33. Breitburg D, Levin LA, Oschlies A, Gregoire M, Chavez FP, Conley DJ, Garcon V, Gilbert D, Gutierrez D, Isensee K et al: **Declining oxygen in the global ocean and coastal waters**. Science 2018, 359(6371):46-+.

34. Hoegh-Guldberg O, Bruno JF: **The impact of climate change on the world’s marine ecosystems**. Science 2010, 328(5985):1523-1528.

35. Hop H, **Gjøsæter H: Polar cod (*Boreogadus saida*) and capelin (*Mallotus villosus*) as key species in marine food webs of the Arctic and the Barents Sea**. Marine Biology Research 2013, 9(9):878–894.

36. Fossheim M, Primicerio R, Johannesen E, Ingvaldsen RB, Aschan MM, Dolgov AV: **Recent warming leads to a rapid borealization of fish communities in the Arctic**. Nature Climate Change 2015, 5(7):673.

37. Holst JC, McDonald A: **FISH-LIFT: a device for sampling live fish with trawls**. Fisheries Research 2000, 48(1):87–91.

38. Tyberghein L, Verbruggen H, Pauly K, Troupin C, Mineur F, De Clerck O: **Bio-ORACLE: a global environmental dataset for marine species distribution modelling**. Global ecology and biogeography 2012, 21(2):272-281.

39. Assis J, Tyberghein L, Bosch S, Verbruggen H, Serrão EA, De Clerck O: **Bio-ORACLE v2. 0: Extending marine data layers for bioclimatic modelling**. Global Ecology and Biogeography 2018, 27(3):277-284.

40. Chabot D, Steffensen JF, Farrell AP: **The determination of standard metabolic rate in fishes**. J Fish Biol 2016, 88(1):81–121.

41. Boutilier RG, Heming TA, Iwama GK: **Physicochemical parameters for use in fish respiratory physiology**. In: Fish physiology. vol. 10: Elsevier; 1984: 403–430.

42. Steffensen J, Bushnell P, Schurmann H: **Oxygen consumption in four species of teleosts from Greenland: no evidence of metabolic cold adaptation**. Polar Biology 1994, 14(1):49–54.

43. Morozov S, McCairns RS, Merilä J: **FishResp: R package and GUI application for analysis of aquatic respirometry data**. Conservation physiology 2019, 7(1):coz003.

44. Scrucca L, Fop M, Murphy TB, Raftery AE: **mclust 5: clustering, classification and density estimation using Gaussian finite mixture models**. The R journal 2016, 8(1):289.

45. Drucker E, Jensen J: **Pectoral fin locomotion in the striped surfperch**. **II. Scaling swimming kinematics and performance at a gait transition**. Journal of Experimental Biology 1996, 199(10):2243–2252.

46. Videler J: **Swimming movements, body structure and propulsion in cod *Gadus morhua***. In: Symp Zool Soc Lond: 1981. 1-27.

47. Brett J: **The respiratory metabolism and swimming performance of young sockeye salmon**. Journal of the Fisheries Board of Canada 1964, 21(5):1183-1226.

48. R Core Team: R: **A language and environment for statistical computing. R Foundation for Statistical Computing**. In.; 2020.

49. Wickham H, Henry L: **tidyr: Tidy Messy Data. R package version 1.1.0**. . 2020.

50. Lindroth A: **Sauerstoffverbrauch der Fische. II. Verschiedene Entwicklungs und Altersstadien vom Lachs und Hecht**. Zeitschrift für vergleichende Physiologie 1942, 29(4):583–594.

51. Ultsch GR, Boschung H, Ross MJ: **Metabolism, critical oxygen tension, and habitat selection in darters (Etheostoma)**. Ecology 1978, 59(1):99–107.

52. Claireaux G, Chabot D: **Responses by fishes to environmental hypoxia: integration through Fry’s concept of aerobic metabolic scope**. Journal of Fish Biology 2016, 88(1):232–251.

53. Neill WH, Bryan JD: **Responses of fish to temperature and oxygen, and response integration through metabolic scope**. Aquaculture and water quality 1991, 3:30–57.

54. Heiss A: **reconPlots: Plot economics graphs with R**. 2022.

55. Fry FEJ: Effects of the environment on animal activity. Pub. **Ontario Fish. Lab. No. 68**. U Toronto Studies, Biol Ser 1947, 55:1–52.

56. Fry F: **The effect of environmental factors on the physiology of fish**. Fish physiology 1971:1–98.

57. Hochachka P: **Scope for survival: a conceptual “mirror” to Fry’s scope for activity**. Transactions of the American Fisheries Society 1990, 119(4):622–628.

58. Seibel BA, Andres A, Birk MA, Burns AL, Shaw CT, Timpe AW, Welsh CJ: **Oxygen supply capacity breathes new life into critical oxygen partial pressure (P_crit_)**. Journal of Experimental Biology 2021, 224(8).

59. Pörtner HO: **Oxygen-and capacity-limitation of thermal tolerance: a matrix for integrating climate-related stressor effects in marine ecosystems**. Journal of Experimental Biology 2010, 213(6):881–893.

60. Rogers NJ, Urbina MA, Reardon EE, McKenzie DJ, Wilson RW: **A new analysis of hypoxia tolerance in fishes using a database of critical oxygen level (P_crit_)**. Conservation physiology 2016, 4(1).

61. Austin MW, Ploughman M, Glynn L, Corbett D: **Aerobic exercise effects on neuroprotection and brain repair following stroke: A systematic review and perspective**. Neurosci Res 2014, 87:8–15.

62. Schurmann H, Steffensen J: **Effects of temperature, hypoxia and activity on the metabolism of juvenile Atlantic cod**. Journal of fish biology 1997, 50(6):1166–1180.

63. Wienerroither R, Johannesen E, Dolgov A, Byrkjedal I, Bjelland O, Drevetnyak K, Eriksen K, Høines Å, Langhelle G, Langøy H: **Atlas of the Barents Sea fishes**. IMR/PINRO Joint Report Series 2011, 1(2011):1-272.

64. Hochachka PW: **Defense strategies against hypoxia and hypothermia**. Science 1986, 231(4735):234–241.

65. Wood SC: **Interactions between hypoxia and hypothermia**. Annual Review of Physiology 1991, 53(1):71–85.

66. Bickler PE, Buck LT: **Hypoxia tolerance in reptiles, amphibians, and fishes: life with variable oxygen availability**. Annu Rev Physiol 2007, 69:145–170.

67. Dunn JF, **Johnston IA: Metabolic constraints on burst-swimming in the Antarctic teleost *Notothenia neglecta***. Marine Biology 1986, 91(4):433–440.

68. Davison W, Forster ME, Franklin CE, Taylor HH: **Recovery from exhausting exercise in an Antarctic fish, Pagothenia borchgrevinki**. Polar biology 1988, 8(3):167–171.

69. Dutil JD, Sylvestre EL, Gamache L, Larocque R, **Guderley H: Burst and coast use, swimming performance and metabolism of Atlantic cod *Gadus morhua* in sub-lethal hypoxic conditions**. Journal of Fish Biology 2007, 71(2):363–375.

70. Chabot D, **Claireaux G: Environmental hypoxia as a metabolic constraint on fish: the case of Atlantic cod, *Gadus morhua***. Marine Pollution Bulletin 2008, 57(6-12):287–294.

71. Wells RM: **Cutaneous oxygen uptake in the Antarctic icequab, *Rhigophila dearborni* (Pisces: Zoarcidae)**. Polar Biology 1986, 5(3):175–179.

72. Hemmingsen EA, Douglas EL: **Respiratory characteristics of the hemoglobin-free fish Chaenocephalus aceratus**. Comparative biochemistry and physiology 1970, 33(4):733–744.

73. Steffensen JF, Lomholt JP, **Johansen K: The relative importance of skin oxygen uptake in the naturally buried plaice, *Pleuronectes platessa*, exposed to graded hypoxia**. Respiration physiology 1981, 44(3):269–275.

74. Hop H, Welch HE, Crawford RE: **Population structure and feeding ecology of Arctic cod schools in the Canadian High Arctic**. In: Fish ecology in Arctic North America American fisheries society symposium: 1997. 68–80.

75. Aune M, Raskhozheva E, Andrade H, Augustine S, Bambulyak A, Camus L, Carroll J, Dolgov AV, Hop H, Moiseev D et al: **Distribution and ecology of polar cod (Boreogadus saida) in the eastern Barents Sea: A review of historical literature**. Mar Environ Res 2021, 166.

76. Welch HE, Crawford RE, **Hop H: Occurrence of Arctic cod (*Boreogadus saida*) schools and their vulnerability to predation in the Canadian High Arctic**. Arctic 1993:331–339.

77. Domenici P, Steffensen JF, Batty R: **The effect of progressive hypoxia on swimming activity and schooling in Atlantic herring**. Journal of Fish Biology 2000, 57(6):1526–1538.

78. Domenici P, Herbert NH, Lefrançois C, Steffensen JF, McKenzie D: **The effect of hypoxia on fish swimming performance and behaviour**. In: Swimming physiology of fish. Springer, Berlin, Heidelberg,; 2013: 129-159.

79. Oppedal F, Dempster T, Stien LH: **Environmental drivers of Atlantic salmon behaviour in sea-cages: A review**. Aquaculture 2011, 311(1-4):1–18.

80. Hochachka PW, Buck LT, Doll CJ, Land SC: **Unifying theory of hypoxia tolerance: Molecular metabolic defense and rescue mechanisms for surviving oxygen lack**. Proceedings of the National Academy of Sciences of the United States of America 1996, 93(18):9493–9498.

81. Divine D, Isaksson E, Martma T, Meijer HAJ, Moore J, Pohjola V, van de Wal RSW, Godtliebsen F: **Thousand years of winter surface air temperature variations in Svalbard and northern Norway reconstructed from ice-core data**. Polar Research 2011, 30.

82. Pavlov AK, Tverberg V, Ivanov BV, Nilsen F, Falk-Petersen S, Granskog MA: **Warming of Atlantic Water in two west Spitsbergen fjords over the last century (**1912**-**2009). Polar Research 2013, 32.

83. Overland J, Dunlea E, Box JE, Corell R, Forsius M, Kattsov V, Olseng MS, Pawlak J, Reiersen LO, Wang MY: **The urgency of Arctic change**. Polar Sci 2019, 21:6–13.

84. Alexander MA, Scott JD, Friedland KD, Mills KE, Nye JA, Pershing AJ, Thomas AC: **Projected sea surface temperatures over the 21st century: Changes in the mean, variability and extremes for large marine ecosystem regions of Northern Oceans**. Elementa-Sci Anthrop 2018, 6.

85. Mark FC, Bock C, Portner HO: **Oxygen-limited thermal tolerance in Antarctic fish investigated by MRI and P-31-MRS**. Am J Physiol-Reg I 2002, 283(5):R1254–R1262.

86. Claireaux G, Lagardère J-P: **Influence of temperature, oxygen and salinity on the metabolism of the European sea bass**. Journal of Sea Research 1999, 42(2):157–168.

87. Halsey LG, Killen SS, Clark TD, Norin T: **Exploring key issues of aerobic scope interpretation in ectotherms: absolute versus factorial**. Reviews in Fish Biology and Fisheries 2018, 28(2):405–415.

88. Brand M, Fischer P: **Species composition and abundance of the shallow water fish community of Kongsfjorden, Svalbard**. Polar Biology 2016, 39(11):2155–2167.

89. Kjesbu OS, Bogstad B, Devine JA, Gjøsæter H, Howell D, Ingvaldsen RB, Nash RD, Skjæraasen JE: **Synergies between climate and management for Atlantic cod fisheries at high latitudes**. Proceedings of the National Academy of Sciences 2014, 111(9):3478–3483.

90. Lapointe D, Guderley H, **Dutil JD: Changes in the condition factor have an impact on metabolic rate and swimming performance relationships in Atlantic cod (*Gadus morhua* L**.). Physiological and Biochemical Zoology 2006, 79(1):109–119.

91. Reidy S, Kerr S, Nelson J: **Aerobic and anaerobic swimming performance of individual Atlantic cod**. Journal of Experimental Biology 2000, 203(2):347–357.

92. Petersen L, **Gamperl A: Effect of acute and chronic hypoxia on the swimming performance, metabolic capacity and cardiac function of Atlantic cod (*Gadus morhua*)**. Journal of Experimental Biology 2010, 213(5):808–819.

93. Fitzgibbon QP, Strawbridge A, Seymour RS: **Metabolic scope, swimming performance and the effects of hypoxia in the mulloway**, Argyrosomus japonicus (Pisces: Sciaenidae). Aquaculture 2007, 270(1-4):358–368.

94. Hop H, **Graham M: Respiration of juvenile Arctic cod (*Boreogadus saida*): effects of acclimation, temperature, and food intake**. Polar Biol 1995, 15(5):359–367.

95. Nahrgang J, Varpe O, Korshunova E, Murzina S, Hallanger IG, Vieweg I, **Berge J: Gender Specific Reproductive Strategies of an Arctic Key Species (*Boreogadus saida*) and Implications of Climate Change**. Plos One 2014, 9(5).

96. Eriksen E, Huserbråten M, Gjøsæter H, Vikebø F, Albretsen J: **Polar cod egg and larval drift patterns in the Svalbard archipelago**. Polar Biology 2019:1–14.

97. Urbanski JA, Litwicka D: **The decline of Svalbard land-fast sea ice extent as a result of climate change**. Oceanologia 2022, 64(3):535–545.

98. Claireaux G, Webber DM, Lagardere JP, **Kerr SR: Influence of water temperature and oxygenation on the aerobic metabolic scope of Atlantic cod (*Gadus morhua*)**. Journal of Sea Research 2000, 44(3-4):257–265.

